# Structural insight into the mechanism of neuraminidase inhibitor-resistant mutations in human-infecting H10N8 Influenza A virus

**DOI:** 10.1101/378075

**Authors:** Babayemi O. Oladejo, Yuhai Bi, Christopher J. Vavricka, Chunrui Li, Yan Chai, Kun Xu, Liqiang Li, Zhe Lu, Jiandong Li, Gary Wong, Sankar Mohan, B. Mario Pinto, Haihai Jiang, Jianxun Qi, George Fu Gao, Po Tien, Yan Wu

**Author notes:** These authors contributed equally to this work (Babayemi O. Oladejo, Yuhai Bi).

## Abstract

The emergence of drug resistance in avian influenza virus (AIV) is a serious concern for public health. Neuraminidase (NA) isolated from a fatal case of avian-origin H10N8 influenza virus infection was found to carry a drug-resistant mutation, NA-Arg292Lys (291 in N8 numbering). In order to understand the full potential of H10N8 drug resistance, the virus was first passaged in the presence of the most commonly used neuraminidase inhibitors (NAIs), oseltamivir and zanamivir. As expected, the Arg292Lys substitution was detected after oseltamivir treatment, however a novel Val116Asp substitution (114 in N8 numbering) was selected by zanamivir treatment. Next generation sequencing (NGS) confirmed that the mutations arose early (after passages 1-3) and became dominant in the presence of the NAI inhibitors. Extensive crystallographic studies revealed that N8-Arg292Lys resistance results mainly from loss of interactions with the inhibitor carboxylate, while rotation of Glu276 was not impaired as observed in the N9-Arg292Lys, a group 2 NA structure. In the case of Val116Asp, the binding mode between oseltamivir and zanamivir is different. Asp151 forms stabilized hydrogen bond to guanidine group of zanamivir, which may compensate the resistance caused by Val116Asp. By contrast, the amino group of oseltamivir is too short to maintain this hydrogen bond, which result in resistant. Moreover, the oseltamivir-zanamivir hybrid inhibitor MS-257 displays higher effectiveness to Val116Asp than oseltamivir, which support this notion.

**Author Summary:** Aside from vaccination, NAIs are currently the only alternative for the clinical treatment and prophylaxis of influenza. Understanding the mechanisms of resistance is critical to guide in drug development. In this study, two drug-resistant NA substitutions, Val116Asp and Arg292Lys, were discovered from oseltamivir and zanamivir treatment of H10N8 virus. Crystal structural analyses revealed two distinct mechanisms of these two resistant mutations and provide the explanation for the difference in susceptibility of different NAIs. Zanamivir and laninamivir were more effective against the resistant variants than oseltamivir, and Arg292Lys results in more serious oseltamivir resistance in N9 than N8 subtype. This study is well-correlated to influenza pandemic/epidemic pre-warning, as the discovery of inhibitor resistant viruses will help for new drug preparedness.

## Introduction

Four NAIs, oseltamivir, zanamivir, peramivir and laninamivir, are currently available for the clinical treatment of influenza virus infections [1–4]. Oseltamivir has been extensively used due to its high oral bioavailability [5], however resistance to both oseltamivir and peramivir is prevalent. On the other hand, although zanamivir and laninamivir offer advantages in terms of drug resistance, both inhibitors are highly polar in their active forms and therefore have lower bioavailability. Furthermore, there are still many known substitutions that lead to zanamivir and laninamivir resistance including Glu119Gly/Ala/Asp, Gln136Lys, Asp151Ala/Asn/Gly/Val, Arg152Lys, Ile222Arg, Asp198Asn, Arg292Lys and Arg371Lys (N2 numbering) [6–10].

Resistance to NAIs usually results from substitutions of highly conserved amino acid residues that form the NA active site. The influenza NA active site contains 8 conserved catalytic residues, and an additional frame of 11 residues that provides structural support to the active site [11, 12]. Substitution of non/semi-conserved residues has also been shown to lead to NAI resistance [13]. Semi-conserved influenza NA residues such as Ile117 and Lys150 have been observed to confer NAI resistance in N1 subtype NA[14, 15], while Gln136 has been observed in both N1 [16] and N2 [17] NA subtypes. However, the mechanisms of drug resistance related to semi-conserved residues are poorly understood [6, 18, 19].

In recent years, some new AIVs, either highly pathogenic (HPAIV) or low pathogen AIV (LPAIV), have emerged with ability to infect humans in addition to H5N1 HPAIV[20–24], due to extensive poultry transportation and wild bird migration [25]. LPAIV H10N8 human-infecting cases were identified in Jiangxi Province, China in 2013 [24, 26]. In December 2013, A/Jiangxi-Donghu/346/2013/H10N8 with the NA-Arg292Lys (N2 numbering, 291 in N8 numbering) mutation was discovered in the trachea aspirate of a 73-year-old patient who died 3 days following oseltamivir treatment [27]. Therefore, H10N8 and other avian-origin influenza A viruses possess a high potential for human pathogenicity and hence pose public health concern.

There are two phylogenetic groups of influenza A NAs (N1, N4, N5 and N8 belong to group 1, while N2, N3, N6, N7 and N9 belong to group 2) [28]. Arg292Lys has been widely reported in group 2 NAs (N2 and N9), while the report of Arg292Lys in a group 1 NAs has never been observed in nature. Structural analysis of N9-Arg292Lys has been shown to result in unfavorable Glu276 conformation for oseltamivir binding [10]. However, the mechanism of Arg292Lys resistance in a group 1 N8 is not well understood due to lack of crystal structures (apo and holo). In this study, we explored the potential drug-resistant substitutions of A/Jiangxi-Donghu/346/2013/H10N8 by passaging the virus in the presence of zanamivir and oseltamivir, respectively. Besides the clinical Arg292Lys substitution which was identified after *in vitro* oseltamivir treatment, a novel Val116Asp substitution was also identified after zanamivir treatment. The residue 116 (residue 114 in N8 numbering) does not play a direct role in active site framework or drug binding and therefore the mechanism underlying Val116Asp resistance is perplexing. We have determined the binding capability (K_m_) of N8 wildtype and two substitutions. The data indicated that Val116Asp and Arg292Lys mutations reduce the affinity between the enzyme and substrate, which results from the conformational shift of the key residues in the active site. Moreover, the inhibition assay demonstrated that the oseltamivir resistance (1,000-fold) caused by Arg292Lys in N8 is much less severe than that of H7N9 (100,000-fold). We have also determined the *apo* crystal structures of N8 wild type and the two mutants and the *holo* structures in complex with oseltamivir, zanamivir, peramivir and laninamivir. These crystal structures reveal the underlying mechanism by which Arg292Lys and Val116Asp mutations develop resistance towards NAIs.

## Results

### In vitro NAI Selective Pressure

H10N8 virus was subjected to 2-fold increases in the concentration of oseltamivir (80-10240 μM) and zanamivir (20-2560 μM) over 8 passages. Prior quantification by plaque assays showed that the virus was resistant to oseltamivir at 13.3 μM and zanamivir at 1.33 μM. After each passage, the hemagglutinin (HA) titer was determined, recorded in triplicate and used to estimate the concentration of virus to be seeded in the next passage. A negative HA titer was observed in the last passage indicating the absence of virus (S1 Table).

### Substitutions under NAI Pressure

Viruses from each passage under inhibitor treatment were subject to high-throughput sequencing for whole genome substitution and intra-cell substitution analysis. The result showed that as expected, the drug-resistant Arg292Lys substitution in NA was detected after oseltamivir treatment; however, a novel NA Val116Asp substitution arose after zanamivir treatment, suggesting a direct role in zanamivir-resistance (S1 Fig). The Arg292Lys mutation under oseltamivir treatment and the Val116Asp mutation under zanamivir treatment arose and became dominant quickly (1-3 passages). Unexpectedly, the pre-exist frequency of Arg292Lys was found to be at 28 *%* in the non-drug treatment control. However, just after 1 passage under oseltamivir treatment, it increased to 92% and levelled off at 92% throughout the next 6 oseltamivir treatment passages (Fig 1A and 2). For zanamivir treatment, the pre-exist frequency of Val116Asp was found to be at 2%; after 1 passage, it increased quickly to 34%, and became absolute dominant (90%) (Fig 1B and 2).

**Fig 1.**
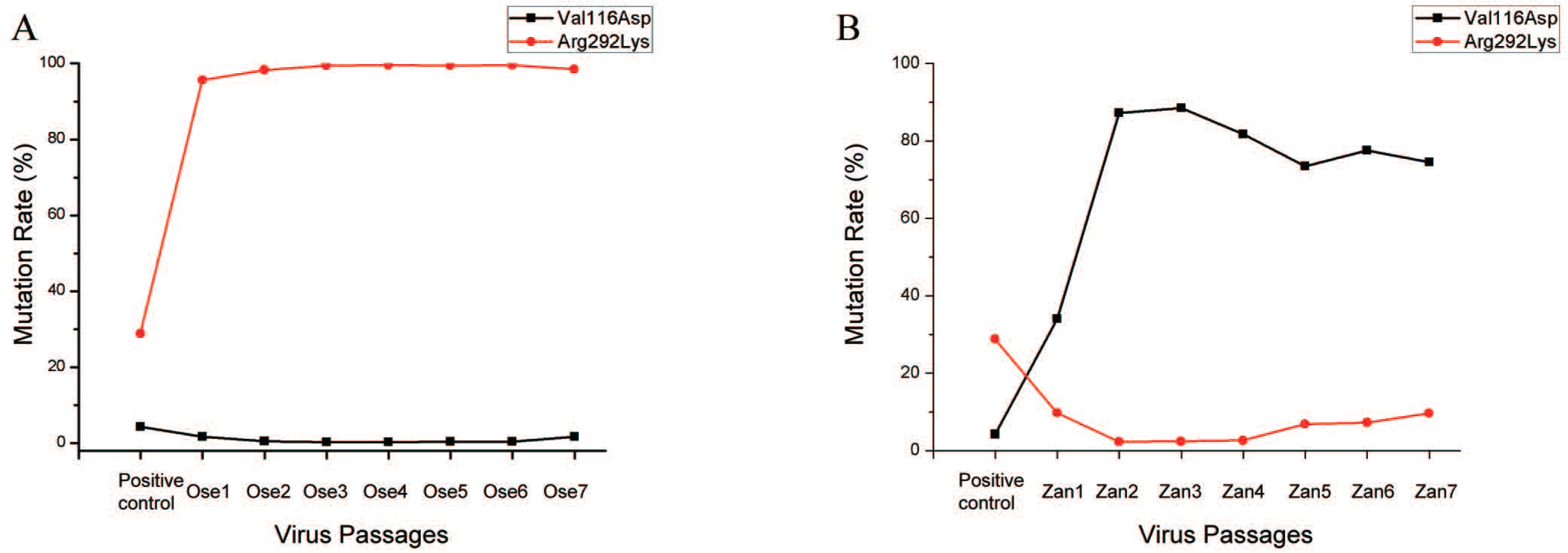
Substitution frequency of Val116Asp and Arg292Lys in N8 under NAI pressure. (A) Mutation frequency of Arg292Lys increased sharply and throughout the passage period during oseltamivir treatment. (B) A similar trend was observed for Val116Asp during zanamivir treatment, although there was a reduction in the mutation rate at passages 5 and 7.

**Fig 2.**
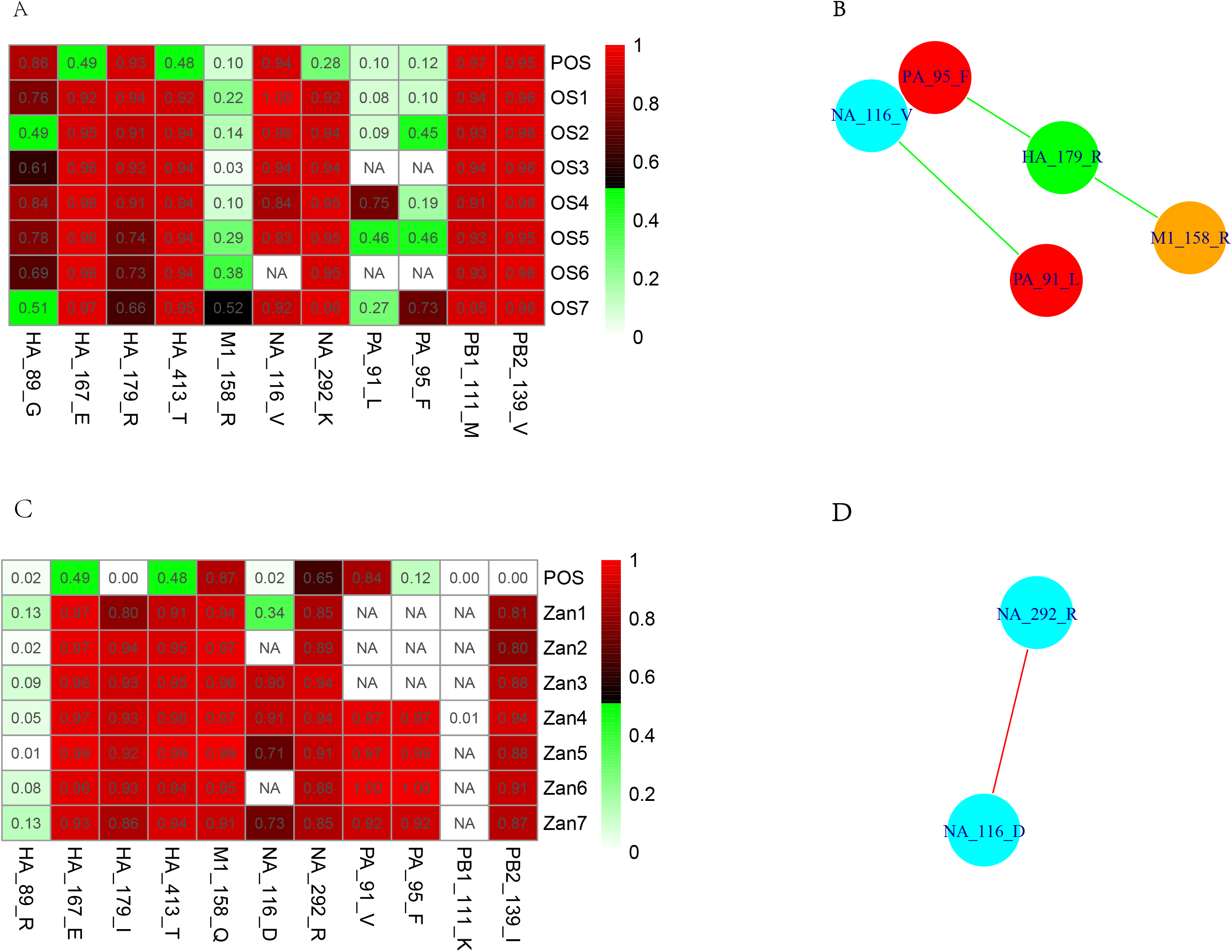
Heatmap and correlation network of oseltamivir treated samples. (A) Heatmap of amino acid residues under oseltamivir treatment from the 1st to 7th passage. (B) Heatmap of amino acid residues under zanamivir treatment from the 1^st^ to 7^th^ passage. The values in each cell represent the substitution frequencies. A sudden change in the default frequency will lead to a change in the amino acid site. The ratio of the depth sequencing site or threshold of total depth is 10. Mutation frequency is based on color change. Each column represents the amino acid sites separated by “-” while each row represents the drug passage including positive control. POS: positive control, N/A: not available data to give a decisive result, OS: oseltamivir, Zan: zanamivir, and the numerals next to each inhibitor represent the passage numbers. (C) Correlation networks of different amino acid sites. (D) Correlation networks of different amino acid sites show a positive correlation between NA Asp116 and NA Arg292. To avoid error, smooth changing sites were used for correlation analysis which could be judged easily from the heatmap. Amino acid sites are indicated as different colors depending on the encoded protein. 2 or more amino acids connected together indicates a correlation larger than 0.8 (r>0.8). Correlations were determined using Pearson’s rank correlation method. P-values of all correlations are less than 0.05. Positive correlation is indicated by a red line connecting amino acids while negative correlation is indicated by a green line.

In addition to these two NA substitutions, several substitutions occurred on the HA and other certain internal genes. Two HA substitutions, Lys167Glu and Ile413Thr, were pre-existing at the frequency of 49% and 48% respectively, and both became dominant under oseltamivir/ zanamivir treatment and kept steady along cell passages with the increase of drug concentration. Importantly, Val139Ile in Polymerase basic protein 2 (PB2) arose and became dominant in the first passage of zanamivir treatment, with the frequency of 81%. However, this was not observed in oseltamivir treatment passages. One common substitution Cys95Phe in Polymerase acidic protein (PA) occurred in both treatment passages. PA Cys95Phe also pre-existed in the drug-free treatment with frequency of 12%, which showed a slow increasing trend and became dominant at passage 7 in oseltamivir treatment. Meanwhile, in zanamivir treatment passages, it became absolute dominant within 3 passages and levelled off. Another PA substitution Val91Leu was only observed in oseltamivir treatment. This mutation pre-existed in the drug-free treatment control at a frequency of 10% and showed a fluctuating trend along passages. Other two substitutions, HA Gly89Arg and M1 Gln158Arg also showed fluctuating trends along oseltamivir treatment passages. Fluctuating trends for HA Gly89Arg mutation was also observed in zanamivir treatment passages (S1 Fig and Fig 2).

Collectively, we can find that most of these substitutions are pre-existing in the original NAI-free cultures. These pre-existing substitutions might be the natural selection pools when under drug treatment. Correlation analysis of these substitutions using their frequencies values found several strong links: one negative link between HA 179Arg and M1 Gln158Arg, one negative link between HA 179Arg and PA Cye95Phe along oseltamivir treatment; one positive link between NA V116Asp and 292Arg. These results suggest that synergy might exist along the dynamic changes of these substitutions, which will be further studied in the future.

### Reduced Substrate Affinity and Inhibition of N8-Arg292Lys and N8-Val116Asp

NA was prepared according to the previously reported methods [29–31]. Lower substrate affinity was observed for N8-Arg292Lys (30-fold) and Val116Asp (6-fold) mutants when compared to the wildtype N8. The recombinant Val116Asp and Arg292Lys variants also exhibited reduced sensitivity to the four clinically used NAIs (oseltamivir, zanamivir, peramivir and laninamivir) relative to the wildtype. The Arg292Lys mutant exhibited higher resistance to all of the four NAIs compared to the Val116Asp mutant. Both mutants showed high resistance to oseltamivir, with a 926-fold and 128-fold increase in mean IC_50_ for Arg292Lys and Val116Asp, respectively. Val116Asp was moderately resistant to zanamivir (40-fold) while Arg292Lys displays much higher zanamivir resistance (704-fold). The resistance pattern of Val116Asp and Arg292Lys to laninamivir is similar to that of zanamivir, with 10-fold and 90-fold lower potencies, respectively. On the other hand, Arg292Lys showed much higher resistance to peramivir with a 3,169-fold increase in mean IC_50_ value, while Val116Asp exhibited a moderate 20-fold reduced sensitivity (Table 1). Interestingly, the oseltamivir-zanamivir hybrid inhibitor, MS-257 was found to be the least affected inhibitor by these mutations when compared to all four clinically used NAIs. The increase in the mean IC_50_ values of MS-257 against Arg292Lys and Val116Asp mutants were 19-fold and 7-fold, respectively.

### Val116Asp and Arg292Lys Result in Subtle N8 Active Site Changes

The crystal structures of native N8, Val116Asp and Arg292Lys were determined at resolutions of 1.9 Å, 1.9 Å and 1.6 Å, respectively (S2 Table). The apo structures of N8 wildtype, Val116Asp and Arg292Lys showed similar overall active site arrangements, with some differences observed regarding Gln136, Thr148, Glu276 and Tyr406 (Fig 3). Specifically, the conformations of 150-loop between N8 wildtype and Val116Asp show slightly different, because of the conformational change of Thr148 and Gln136. The conformations of 430-loop between N8 wildtype and Arg292Lys display variable configurations. Moreover, the side chain of Gln136, Arg118, Tyr406 and Glu276 between these two native structures exhibit different conformations. All these observed conformational differences explain why the two N8 mutants have lower affinity to the substrate compared to N8 wildtype.

**Fig 3.**
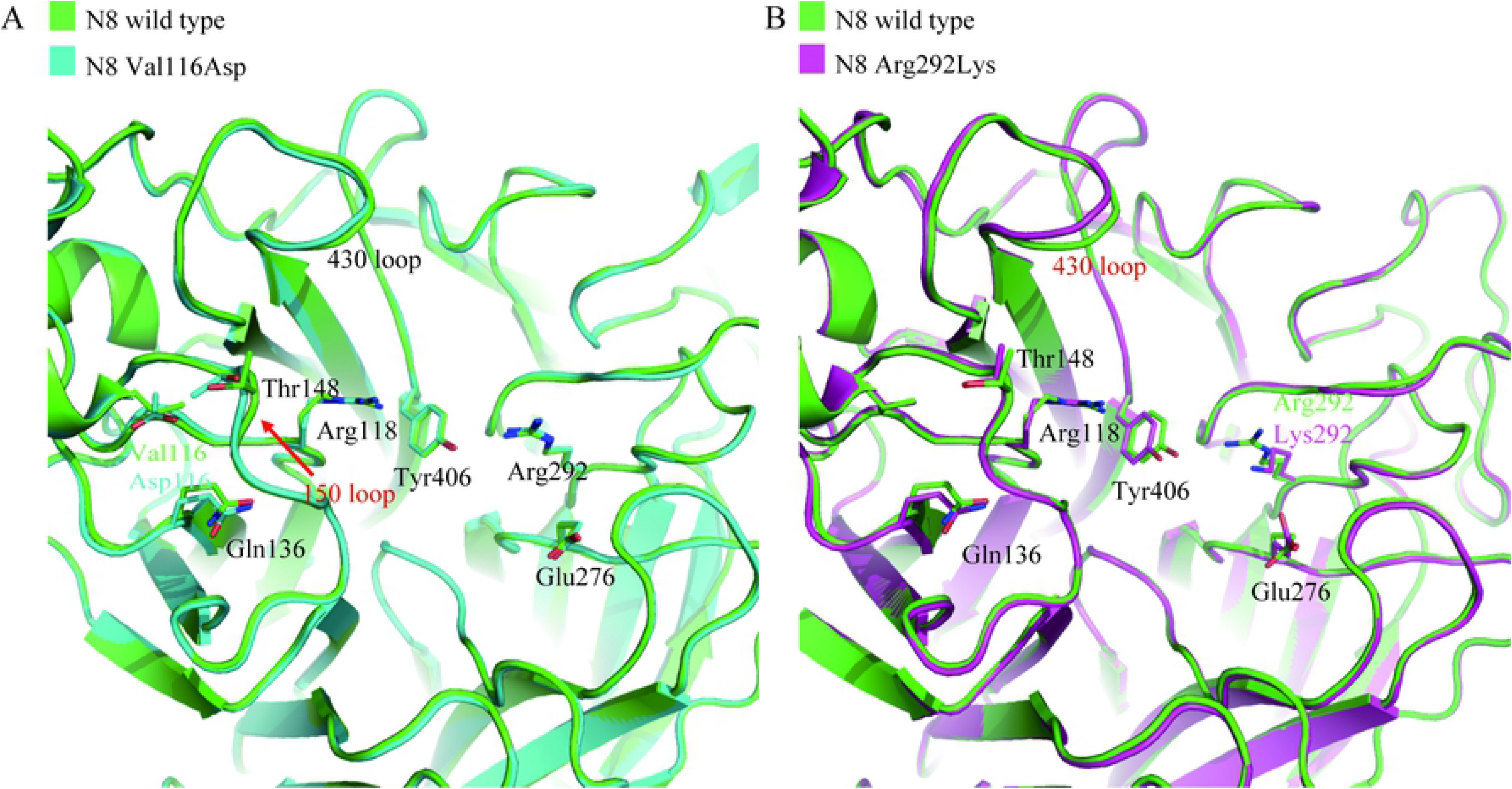
Active site comparison of N8 wildtype (green), Val116Asp (cyan) and Arg292Lys (magenta). (A) The comparison of the key residues conformation between N8 wildtype and N8-Val116Asp. (B) Comparison of the key residues in the active site between N8 wildtype and N8-Arg292Lys. Key residues are labeled in sticks.

### A Distinct Mechanism of Arg292Lys Drug Resistance in Group 1 N8

To understand the mechanisms of N8 drug resistance, inhibitor complex structures with wildtype or mutant N8 were compared. The crystal structures of wildtype N8 complexed with zanamivir, oseltamivir, laninamivir and peramivir were determined at resolutions of 1.8 Å, 2.1 Å, 1.8 Å and 2.0 Å, respectively (S3 Table). N8-Arg292Lys complexes were determined at resolutions of 1.9 Å, 1.8 Å, 2.0 Å and 1.8 Å for zanamivir, oseltamivir, laninamivir and peramivir, respectively (S4 Table). Binding of oseltamivir, zanamivir, peramivir and laninamivir to wildtype H10N8 NA resembles that of typical group 1 NA binding (Fig 4A, C, E and G), with differences observed regarding Tyr347 in both oseltamivir and laninamivir complex structures. In the wildtype N8 complexes with zanamivir, oseltamivir and laninamivir, as well as the Arg292Lys complex with zanamivir, Tyr347 hydrogen bonds with Arg371 and the inhibitor carboxylate. In the wildtype peramivir complex, as well as the Arg292Lys complexes with laninamivir, oseltamivir and peramivir, Tyr347 points away from the active site where it can no longer interact with the inhibitor.

**Fig 4.**
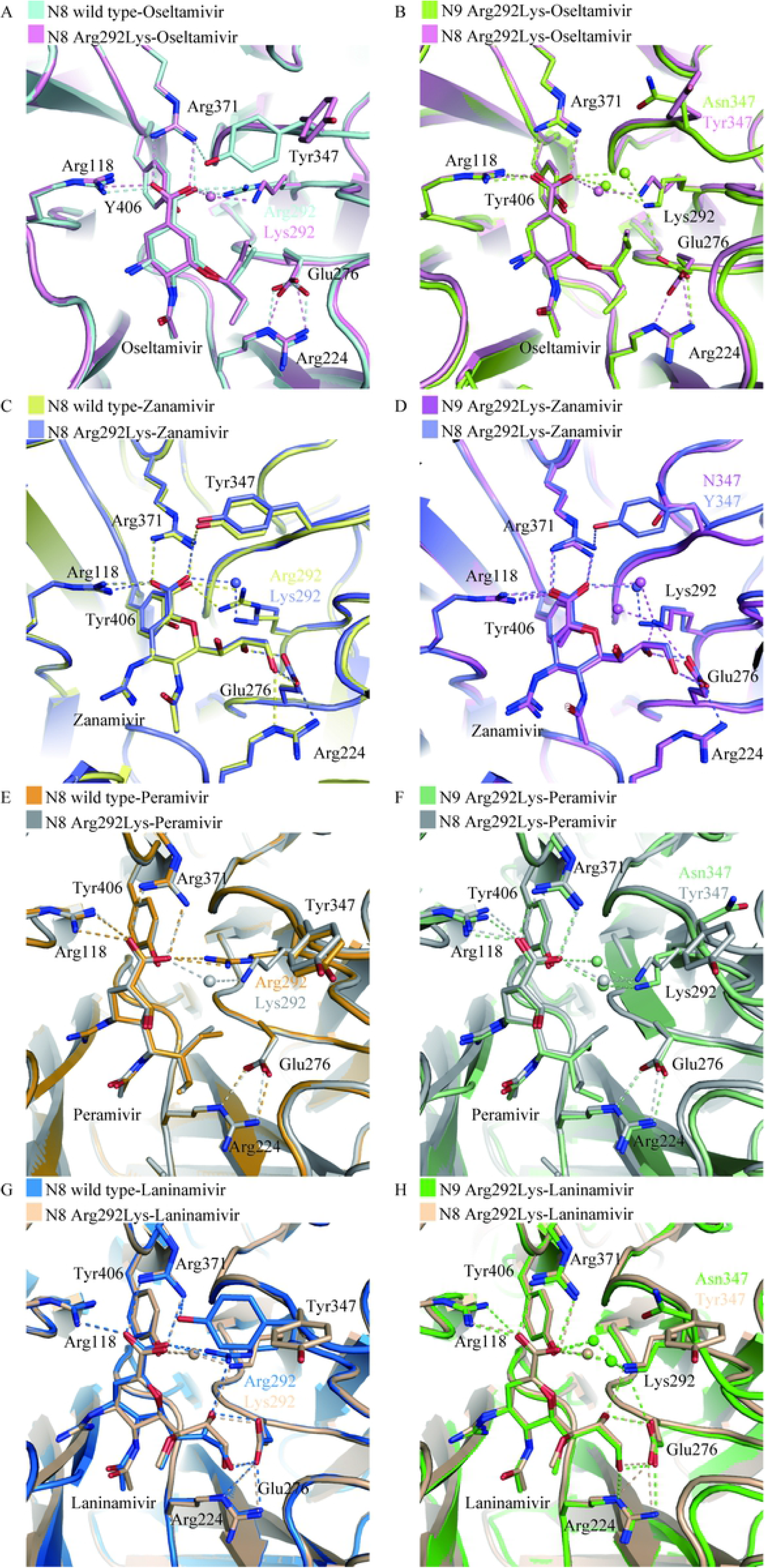
Comparison of wildtype N8, N8-Arg292Lys and N9Arg292Lys binding to four NAIs. (A and B) oseltamivir; (C and D) zanamivir; (E and F) peramivir and (G and H) laninamivir. All the inhibitors and the key residues are displayed in sticks. The hydrogen bonds are indicated in dash. The PDB code of N9-Arg292Lys-oseltamivir, N9-Arg292Lys-zanamivir, N9-Arg292Lys-peramivir and N9-Arg292Lys-laninamivir are 4MWW, 4MWX, 4MX0 and 4MWY, respectively.

Arg292 is part of the NA tri-arginine cluster that forms strong ionic interactions with the first-generation NAI carboxylates. In the structures of N8-Arg292Lys complexed with the four NAIs, Lys292 interacts with the carboxylate group of NAIs by a bridging water molecule (Fig 4A, C, E and G). The Glu276 adopts similar conformation in N8-Arg292Lys-zanamivir and laninamivir complexes (Fig 4C and G), but slightly differ in oseltamivir and peramivir-complexes (Fig 4A and E).

The binding of oseltamivir to N8 (group 1 NA) and N9 (group 2 NA) was carefully compared (Fig 4B, D, F and H). The flexibility of residue Glu276 is observed in the N9-Arg292Lys oseltamivir complex structure, but only a slightly shift in the N8-Arg292Lys oseltamivir complex structure. Specifically, in N8-Arg292Lys, Glu276 rotates towards Arg224 for optimal oseltamivir binding, which results in a mild shift of the oseltamivir hydrophobic group (1.11 Å). In contrast, the side chain of Glu276 in N9-Arg292Lys is oriented toward the oseltamivir carboxylate hydrophobic pentyloxy group, which is pushed 2.98 Å away from the active site. Furthermore, Arg292Lys substitution in N8 has no effect on the hydrogen bonds between Glu276 and Arg224, while, in N9, this substitution results in loss of one hydrogen bond between Glu276 and Arg224 (Fig 4B). These differences help to explain the observation that the effect of N8-Arg292Lysoseltamivir resistance is 1000-fold whereas the effect of N9-Arg292Lys resistance is 100,000-fold [10].

Notably, in the structure of N8-Arg292Lys-zanamivir, the orientation of Tyr347 is the same as N8 wildtype, which forms hydrogen bond with the side chain of Arg371 (Fig 4C). However, the residue 347 in N9 is Aspartic acid, which side chain is not enough to form hydrogen bond with Arg371 (Fig 4D). Therefore, zanamivir shows better inhibition to N8 than N9 with Arg292Lys substitution (Table 1).

### The guanidine group of zanamivir compensates the Val116Asp resistant N8 substitution

In order to understand the mechanism of Val116Asp resulting in more severe resistance to oseltamivir than zanamivir, the zanamivir and oseltamivir complex structures of N8-Val116Asp were determined at resolutions of 1.9 Å and 2.1 Å, respectively (S5 Table). The 150-loop of inhibitor bound-N8 (Val116Asp) complexes exhibited closed conformation, in which the Asp151 interacts with inhibitors. When compared the oseltamivir complex-N8 wild type with that of the Val116Asp mutant, it was found that the side chain of Asp151 in N8 wildtype hydrogen bonds to C4-amino group of oseltamivir with a distance of 2.70 Å, while the corresponding distance in N8-Val116Asp was found to be 3.34 Å. In the case of zanamivir complexed structures, oxygen atom on the main chain of Asp151 makes hydrogen bond interactions with the guanidine group of zanamivir, with distance of 2.88 Å and 3.03 Å in N8 wildtype and N8-Val116Asp mutant, respectively. The hydrogen bond interactions of zanamivir guanidine group withAsp151 is less affected by the Val116Asp substitution than that of oseltamivir amino group. Moreover, Tyr347 in these two Val116Asp complex structures are shifted away from both inhibitors, while Tyr347 hydrogen bonds with the zanamivir and oseltamivir carboxylates groups in wildtype N8 complex structures (Fig 5). Interestingly, the oseltamivir-zanamivir hybrid inhibitor MS-257 showed better inhibition to N8-Val116Asp than oseltamivir (Table1).

**Fig 5.**
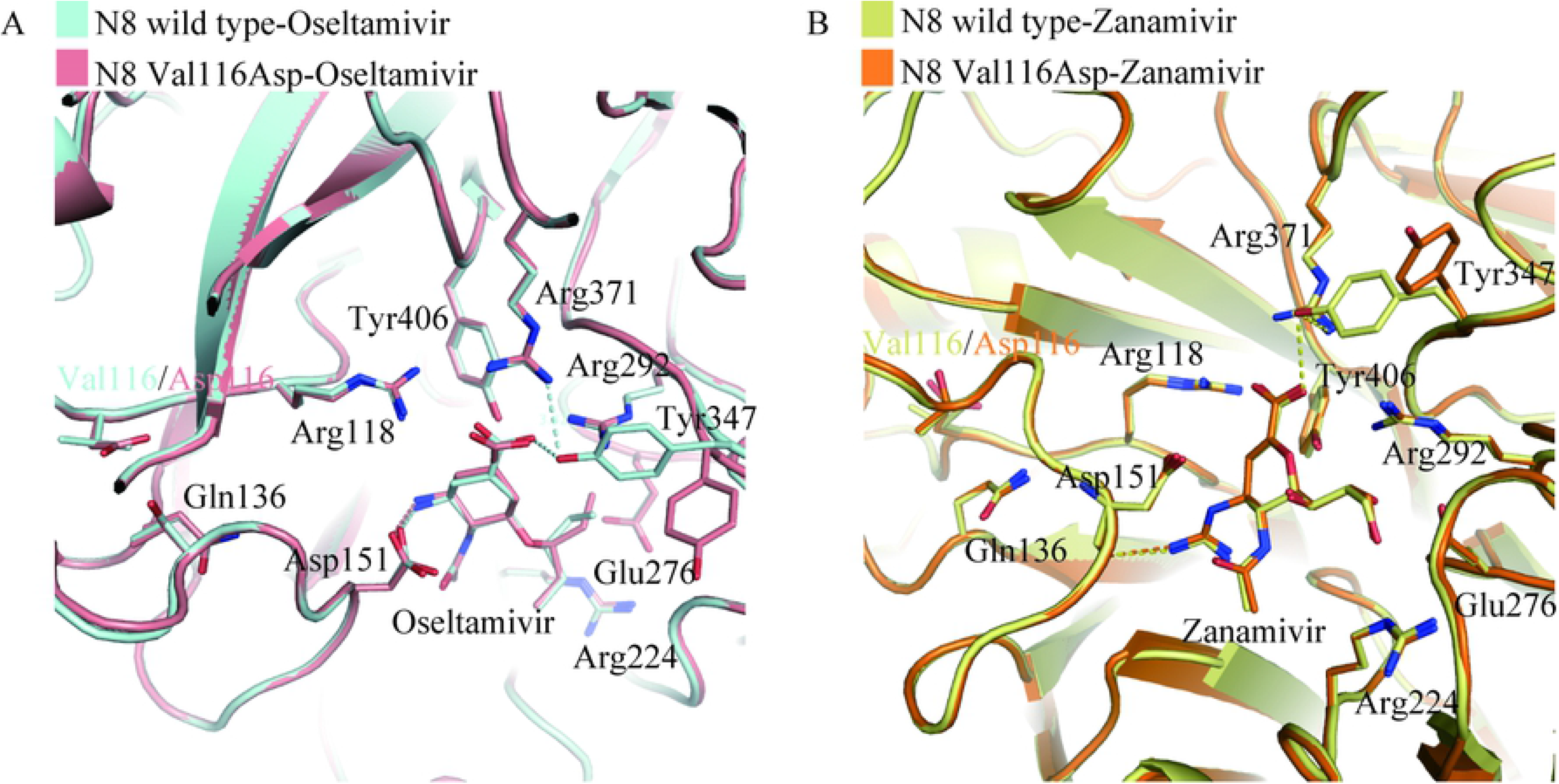
Comparison of the binding mode between N8 wildtype and N8-Val116Asp with two inhibitors. (A) The binding mode of N8 wildtype-oseltamivir (cyan) and N8-Val116Asp-zanamivir (salmon). (B) The binding mode of N8 wildtype-zanamivir (bright yellow) and N8-Val116Asp-zanamivir (orange).

## Discussion

Although the crystal structures of all 9 influenza A NA subtypes have been solved to date [10, 28, 30–37], the influenza A virus is constantly adapting and many new variations are being discovered, especially those with drug resistance. For example, the crystal structure analysis of Arg292Lys mutant of group 2 N9 was completed in 1998 [38], yet the corresponding group 1 N8 structure reported here contains unique features which are clearly observed during oseltamivir binding. These structural differences are also reflected in the K_m_ values which increased 30-fold for N8-Arg292Lys and 88-fold for H7N9 N9-Arg292Lys [10], relative to the corresponding wildtype NAs.

The Val116Lys mutation was the most intriguing N8 substitution. It is challenging to predict a precise mechanism for NAI resistance, because the site of mutation is distal from the active site frame work. Prior to solving the crystal structures, we anticipated that the Val116Asp substitution might affect the interaction between Arg118 and the carboxylate group of the inhibitor, however contrary to expectation, the N8-Val116Lys complex structures clearly suggested that this is not the case and the loss of Tyr347-inhibitor interaction as the resistance mechanism. Tyr347 is found only in group 1 NAs which should contain the 150-loop cavity [28]. This residue was also observed to be a key factor in explaining the slightly higher NAI inhibition observed in N5 (typical group 1 NA) [36] relative to pandemic 09N1 (atypical group 1 NA) [39] and 57N2 (group 2 NA) [31]. Thus, we previously speculated that Tyr347 might compensate for the open 150-loop in regards to substrate binding [31].

Val116Lys also resulted in an altered conformation of the 150-loop residue Thr148 (Fig 3). It is possible that the further changes observed in the loop residues 146-148 upon oseltamivir binding might affect the ionic interaction of Asp151 with the oseltamivir amino group (Fig 5). Despite the moderate effect on NAI inhibition, the Val116Asp substitution also resulted in a 6-fold K_m_ increase relative to the wildtype (Table 1). This indicates that Val116Asp also interferes with substrate binding.

Zanamivir and laninamivir are more similar to the human sialic acid, N-acetylneuraminic acid, than oseltamivir and peramivir, and therefore should be less susceptible to drug-resistance. Of the 4 clinical NAIs, zanamivir bound to both mutants in the most optimal conformation, whereas oseltamivir binding was the least optimal. Resistance of both N8 variants to zanamivir and laninamivir was also lower than that of oseltamivir and peramivir.

Some questions remain from the present analysis, including why peramivir was the most potent inhibitor of wildtype N8 despite lacking any Tyr347 interactions. Moreover, binding of the prodrug laninamivir octanoate (CS-8958) to N8 was distinct from group 1 09N1 and similar to group 2 N2, which indicates that NAI binding is not always group specific.

In summary, our current study revealed the 4-clinical available NAI resistant substitutions for N8 and the underlying mechanisms have also been structurally delineated, which will help for next-generation drug development guide for drug usage in future prewarning of H10N8 AIV human infections.

## Materials and Methods

### Cells, Virus and NAIs

A549 cells were obtained from China Infrastructure of Cell Line Resources, Beijing and were grown in Dulbecco Modified Eagle’s Medium (DMEM) (Gibco by Life Technologies Incorporation™, Grand Island, New York, USA) containing 5% fetal bovine sera (FBS) (Irvine Scientific). The human influenza A/Jiangxi-Donghu/346/2013(H10N8) virus of avian origin was propagated in the allantoic cavity of 10-day-old fertilized specific-pathogen-free (SPF) hen eggs (Beijing Vital River Laboratory Animal Technology Co., Ltd.) for 96 hours at 37°C. A hemagglutination assay was carried out on the harvested infected allantoic fluid in 96-well plates. After adding 25 μL of PBS to each well, 25 μL of virus suspension was added to the first well, followed by a series of 2-fold dilutions and gentle mixing. Next, about 25 μL of freshly prepared 1% chicken RBC was added to each well and left for 30 minutes at room temperature. Clear hemagglutination of the allantoic fluid containing the virus with chicken RBC indicates viral growth and confirms the presence of the virus. Influenza neuraminidase inhibitors oseltamivir acid (GS 4071) and zanamivir were synthesized at MedChem Express LLC, NJ, USA. MS-257 was provided by cooperator Prof. Mario Pinto.

### Isolation of H10N8 Variants with Decreased Susceptibility to Oseltamivir and Zanamivir

24-hour-old A549 cells were grown in 24 well tissue culture plates and infected with egg-grown virus at a low multiplicity of infection (MOI 0.001 PFU per cell) in maintenance medium. The virus was allowed to adsorb for only 15 minutes at 37°C, after which the inoculum was removed, and the cells were washed twice with pre-warmed PBS followed by addition of 1 ml of each drug preparation to the seeded infected cells. The plates were incubated at 37°C for 72 hours. The virus titer in each culture supernatant was determined by hemagglutination of chicken RBC. The culture supernatant that contained the minimal dose drug that still resulted in cytopathy and detectable HA titer was used as inoculum to infect new cell monolayers at a low MOI. The virus from that sample was then allowed to grow in the presence of a series of 2-fold oseltamivir and zanamivir dilutions. The virus concentrations used in the selection protocol varied, depending on the HA titer in the preceding passage and about 1 ml aliquot of viral stock culture supernatant from the preceding passage was used to infect the fresh cell monolayer cells. The HA titer was determined after each passage to estimate the amount of virus needed for infection. This selection was carried out for a total of 8 passages for both inhibitors in triplicates. Drug concentration ranged from 80 μM to 10.24 mM for oseltamivir and 20 μM to 2.56 mM for zanamivir. All conditions were carried out in triplicates.

### Virus RNA Extraction and Next-Generation Sequencing (NGS)

RNA was extracted from virus isolates using a QIAamp viral RNA Mini Kit from QIAGEN, Germany (Cat. No. 52904). Complete influenza A genomes were prepared from the RNA using the Takara PrimeScriptTM One step RT-PCR Kit Ver. 2 (TAKARA BIO INC. Cat. #RR055A v201309Da, Japan) with slight modifications. Each RNA preparation was mixed with the enzyme mix in a 50 μL reaction system. The thermal cycle parameters were 50°C for 30 mins, 94°C for 2 mins, and then 35 cycles at 94°C for 30 secs, 58°C for 30 secs, and 72°C for 3 mins 20 secs. Primers used were 20 μM MBTuni-12 (5’-ACGCGTGATCAGCAAAAGCAGG) and MBTuni-13 (5’-ACGCGTGATCAGTAGAAACAAGG) that correspond to the 5’ and 3’ conserved sequences of all eight influenza A segments (26). NGS was used to determine the whole genomes of treated H10N8 samples. The sequencing libraries were prepared from H10N8 whole genome PCR products by end-repairing, dA-tailing, adapter ligation and PCR amplification, according to the manufacturer instructions (Life technologies). The libraries were sequenced on an Ion Proton™ System, and sequencing depth was 1 G base per sample. After sequencing, raw NGS short reads were processed by filtering out low-quality reads, adaptor-contaminated reads (with >15 bp matched to the adapter sequence), poly-Ns (with 8Ns) (SOAP2 (v2.21), <5 mismatches). Clean short reads were then mapped onto the eight reference genome segments (A/Jiangxi-Donghu/346/2013(H10N8)) using TMAP (v3.4.1) (https://github.com/iontorrent/TS/tree/master/Analysis/TMAP) with a match rate larger than 0.90, producing the short-read-reference-genome mapping file (SRRG file), and the dominant base on each site was called to obtain the consensus genome. Consensus sequence alignments were done using MUSCLE [40] to identify the variable sites. The consensus nucleotide sites, which have changed in more than any one sample comparing with the original genome (A/Jiangxi-Donghu/346/2013 (H10N8)), are designated as variable sites. Intra-host single nucleotide variations (iSNV) and their correlations were analyzed based on the SRRG file.

### Expression and Purification of Influenza Virus NA

Recombinant NA protein was prepared using the established baculovirus expression system (19, 20, 22). The ectodomain (residues 81 to 469 in N8 numbering) of A/JD/346/2013/H10N8 was cloned into pFastBac1 baculovirus transfer vector (Invitrogen) and the Val114Asp and Arg291Lys substitutions were constructed by site-directed mutagenesis PCR based on N8 wildtype and expressed in a baculovirus system for structural and functional analysis. A GP67 signal peptide was added at the N terminus to facilitate secretion of the recombinant protein, followed by a His tag, a tetramerizing sequence, and a thrombin cleavage site. Recombinant pFastBac1 plasmid was used to transform DH_10_Bac™ *Escherichia coli* (Invitrogen). The recombinant baculovirus was obtained following the manufacturer’s protocol, and Hi5 cell suspension cultures were infected with high-titer recombinant baculovirus. After growth of the infected Hi5 suspension cultures for 2 days, centrifuged media were applied to a 5-mL HisTrap FF column (GE Healthcare), which was washed with 20 mM imidazole. The NA was thereafter eluted using 300 mM imidazole. After dialysis and thrombin digestion (3 U/mg NA; BD Biosciences) overnight at 4°C, gel filtration chromatography was performed with a Superdex^®^ 200 10/300 GL column (GE Healthcare) using 20 mM Tris-HCl and 50 mM NaCl (pH 8.0) buffer or PBS buffer for crystallization or functional assay, respectively. Pure NA fractions were selected and further concentrated using a 10 kDa (Millipore) membrane concentrator.

### Crystallization, Drug-Soaking and Crystal Structure Determination

wildtype and mutant N8 crystals were obtained using the sitting-drop vapor diffusion method. The NA proteins [1 μL of10mg/mL protein in 20mMTris and 50mMNaCl (pH 8.0)] were mixed with 1 μL of reservoir solution. N8 wildtype crystals were obtained in the condition of 0.1 M sodium acetate trihydrate pH 4.7 and 5% w/v Polyethylene glycol 10,000. N8-Val116Asp crystals were obtained in the condition of 0.1M DL-malic acid pH 7.0 and 12% Polyethylene glycol 3,350. N8-Arg292Lys crystals were obtained in the condition of 0.1 M potassium phosphate monobasic/sodium phosphate dibasic pH 6.2 and 10% Polyethylene glycol, 3,000.

All NA crystals were cryoprotected in mother liquor with the addition of 20% (vol/vol) glycerol before being flash-cooled at 100 K for obtaining apo structures. The crystals were then incubated in the mother liquor containing 20 mM inhibitors (oseltamivir acid, zanamivir, peramivir, and laninamivir) and then flash-cooled at100 K.

Diffraction data were collected at Shanghai Synchrotron Radiation Facility beamline BL17U. The collected intensities were indexed, integrated, corrected for absorption, and scaled and merged using HKL-2000 (27). The structure of N8 was solved by molecular replacement using Phaser (28) from the CCP4 program suite (29), with the structure of N8 (PDB ID code 2HT7) as a search model. N8-Val116Asp and Arg292Lys were equally solved using the native N8 as the search model. The initial model was refined by rigid body refinement using REFMAC5 (30), and extensive model building was performed using COOT (31). Further rounds of refinement were performed using the phenix.refine program implemented in the PHENIX package (32) with energy minimization, isotropic ADP refinement and bulk solvent modeling. The stereochemical quality of the final model was assessed with PROCHECK (33). Structures were aligned and analyzed using PyMOL.

### Fluorescent NA Activity Assay

A 4-methylumbelliferyl-Neu5ac-(MUNANA)-based fluorometric NA assay (34) was used for measuring the NA activity and inhibition. The appropriate protein concentrations were chosen after several rounds of preliminary tests. The km values for the active NAs were determined by mixing 10 μL of each recombinant protein with 10 μL of buffer MES-CaCl2 buffer (pH 6.0) in each 96-well plate and serial dilutions (9.76 μM - 5mM) of MUNANA (30 μL) were added to each well. The fluorescence intensity of the released product was measured every 30 seconds for 1 hour at 37°C on the SpectraMax M5 (Molecular Devices), with excitation and emission wavelengths of 355 nM and 460 nM, repectively. For inhibition assays, 10 μL of recombinant protein was mixed with 10 μL of PBS or inhibitor in 96-well standard opaque plate and 30 μL of 167 μM MUNANA (Sigma, USA) in 33 mM MES and 4 mM CaCl2 (pH 6.0) for a final substrate concentration of 100 μM. The inhibitors (in different concentrations) were pre-incubated with the NA protein for 30 minutes at 37°C before adding MUNANA, and then loaded on the SpectraMax^®^ M5 Molecular devices. Fluorescence was monitored immediately after substrate addition at 1-minute intervals for 30 minutes. All assays were performed in triplicates and IC50 values for each inhibitor were calculated with Graphpad Prism 5.0 as the concentration of inhibitor resulting in a 50% reduction in fluorescence units (FU) compared with the control.

## Supporting Information Legends

**Table 1.** K_m_ and IC_50_ values for N8 wildtype and mutant proteins. The IC_50_ values and 95% confidence intervals (CIs) are provided.

**S1 Fig.** Consensus sequence variations of H10N8 virus under drug treatment. The first row represents the structural proteins of H10N8 and vertical numbers represent different amino acid sites. The consensus amino sites list default amino acids, dots represent no substitutions and question marks represents unknown amino acids due to sites that were not covered by short reads. Osel: oseltamivir, Zan: zanamivir, and the numeral to the right of Osel and Zan are the passage numbers.

**S1 Table.** Inhibitor concentration and HA titer of passaged viruses. The inhibitor concentration and HA titer for eight passages are listed in the table. “-” means no hemagglutination.

**S2 Table.** Crystallographic data collection and refinement statistics of native N8 and N8 mutations.

**S3 Table.** Crystallographic data collection and refinement statistics of N8-inhibitor complexes.

**S4 Table.** Crystallographic data collection and refinement statistics of N8-Arg292Lys–inhibitor complexes.

**S5 Table.** Crystallographic data collection and refinement statistics of N8-Val116Asp–inhibitor complexes

